# Amino acid availability is not essential for lifespan extension by dietary restriction in the fly

**DOI:** 10.1101/2021.07.10.451902

**Authors:** Sarah L Gautrey, Mirre J P Simons

## Abstract

Dietary restriction (DR) is one of the most potent ways to extend health- and lifespan. Key progress in understanding the mechanisms of DR, and ageing more generally, was made when dietary protein, and more specifically essential amino acids (EAA), were identified as the key dietary component to restrict to obtain DR’s health and lifespan benefits. This role of dietary amino acids has strongly influenced work on ageing mechanisms, especially in nutrient sensing, e.g. Tor and insulin(-like) signalling networks. Experimental biology in *Drosophila melanogaster* has been instrumental in generating and confirming the now dominant hypothesis that EAA availability is central to ageing. Here, we expand on previous work testing the involvement of EAA in DR through large scale (N=6,238) supplementation experiments across four diets and two genotypes in female flies. Surprisingly, we find that EAA are not essential to DR’s lifespan benefits. Importantly, we do identify the fecundity benefits of EAA supplementation suggesting the supplemented EAA were bioavailable. Furthermore, we find that the effects of amino acids on lifespan vary by diet and genetic line studied and that at our most restricted diet fecundity is constrained by other nutrients than EAA. We suggest that DR for optimal health is a concert of nutritional effects, orchestrated by genetic, diet and environmental interactions. Our results question the universal importance of amino acid availability in the biology of ageing and DR.

## Introduction

A key step in understanding the mechanisms underlying dietary restriction (DR) is to identify which components of the diet cause increased longevity and decreased fecundity, and whether different nutrients are responsible for these related but potentially separate responses^1–3^. When the precise nutrients that cause the health benefits of DR are understood, it is probable more precise molecular mechanisms of DR can be distilled. In addition, the health benefits of DR can potentially be separated from its side-effects such as those on reproduction, or it could be that these effects cannot be separated as they are so physiologically entwined^3–5^.

Despite considerable research we still do not know the exact nutrient, or combination of nutrients, responsible for lifespan extension seen under DR^1,3,6,7^. Even for one of the most tractable and studied model organisms in this regard, the fruit fly (*Drosophila melanogaster*), the precise effects of separate dietary components are far from fully elucidated^8–12^. A consistent and growing line of research has, however, identified protein restriction as the causative route for lifespan extension in flies^10,13–15^, and a wide range of other species, e.g. crickets^16^, stickleback^17^ and mice^18^. These experiments using the Geometric Framework, in which the ratio and amounts of macronutrients in semi- or fully defined diets are varied, demonstrated that the macronutrient protein is the main dietary axis determining lifespan. In flies, in which this paradigm has been most rigorously applied, experimental artefacts curtailing lifespan due to capillary feeding have been suggested. Yet, food delivery via vials, void of these artefacts, yielded similar conclusions^19,20^.

The most conclusive evidence for protein as the main dietary component determining lifespan comes from supplementation studies of protein to a DR diet (in the form of casein) which nullified DR’s longevity effect^13,21^. One of the most influential studies in this field used a range of supplementation studies to pinpoint restriction of essential amino acids (EAA) as responsible for the DR^5 replicated in 22^. Supplementing the DR diet with EAAs led to an increase in fecundity and decrease in lifespan phenocopying the fully fed diet. Of these EAA, Methionine alone was shown to determine the fecundity response to DR, with lifespan being determined by a combination of Methionine and other EAA. This insight fits with work on rodents, in which restriction of Methionine can extend lifespan^23,24^. In flies, restriction of Methionine extends lifespan, but only under conditions of low amino acid status^25^. In mice, EAA supplementation nullified DR’s lifespan extension^26^ and restriction of particular amino-acids, namely branched-chained, can confer health but not necessarily lifespan benefits^1,7,27^.

The key role of EAA in DR further fits with the identification of Tor and IGF as central cellular signalling pathways in the molecular biology of ageing^28^, as these pathways sense nutrient availability^1,29^. There is thus a central line of literature within the biology of ageing field that implicates amino acid availability as a key determinant of ageing^30,31^. This accepted concept has strongly influenced how we view the mechanisms of ageing more generally. Although, in the fly, surprisingly, DR has additive benefits to lifespan on top of those gained by Tor reduction^32^ and reduced insulin-like signalling^33^. Across model organisms, it has been suggested Tor, somatotrophic signalling and DR in general might act through different physiology to extend lifespan^34,35^. Recent insights implicating other nutrients in the DR reproduction and longevity response^8^, the emerging criticism of the importance of dietary protein over calories in mice^36^ and the demonstrated independent effects of sugar^11,37,38^ whilst largely absent in the geometric framework^14^, have now cast doubt on the universal importance of EAA in DR.

Here we report on an EAA supplementation experiment, using four different diets (varying in carbohydrate to protein ratio) and two different genotypes. We find that adding EAA had little effect on lifespan. Furthermore, we find that the modulation of lifespan by EAA depend on which diet it is supplemented to (note, previous work exclusively supplemented DR diets) and on the genetic line studied. In contrast, we did find the expected increase in fecundity when EAAs were supplemented, and these effects were similar across the two genetic lines studied. Our results question the universal importance of EAA in DR, by revisiting and expanding prior influential work on the fly in this area, thereby identifying a potential dogma in the biology of ageing field.

## Results & Discussion

The effects on lifespan of EAA addition were dependent on the genetic line tested and on the specific diet it was supplemented to (significant three-way interaction between diet, line and EAA addition, χ2=11.8, df=4, P=0.019, N=6,238). Within the *yw* genotype the direction and magnitude of the effect of EAA varied depending on the diet which it is supplemented to (interaction: χ2=10.9, df=3, P=0.012, N=3,103). The overall effect on lifespan of amino-acid addition was however only a fraction of that induced by diet, a log hazard of 0.19, compared to hazard differentials of −1.35 and 1.08 when yeast is manipulated in the diet (Table S1). Within genotype *195* the effect of amino-acid addition did not vary significantly with the diet it was supplemented to (χ2=3.45, df=3, P=0.33, N=3,135). Again, the effect of EAA on log hazard was modest (0.31±0.13, P=0.015) in comparison to the effects induced by varying yeast concentration (−0.66; 2.76, Table S1). It is evident therefore that the effects of EAA addition did not recapitulate the effects of DR observed when varying yeast concentration in the diet (Figure 1), suggesting EAA are at best only partially responsible for the DR longevity responses we observed.

**Figure 1.**
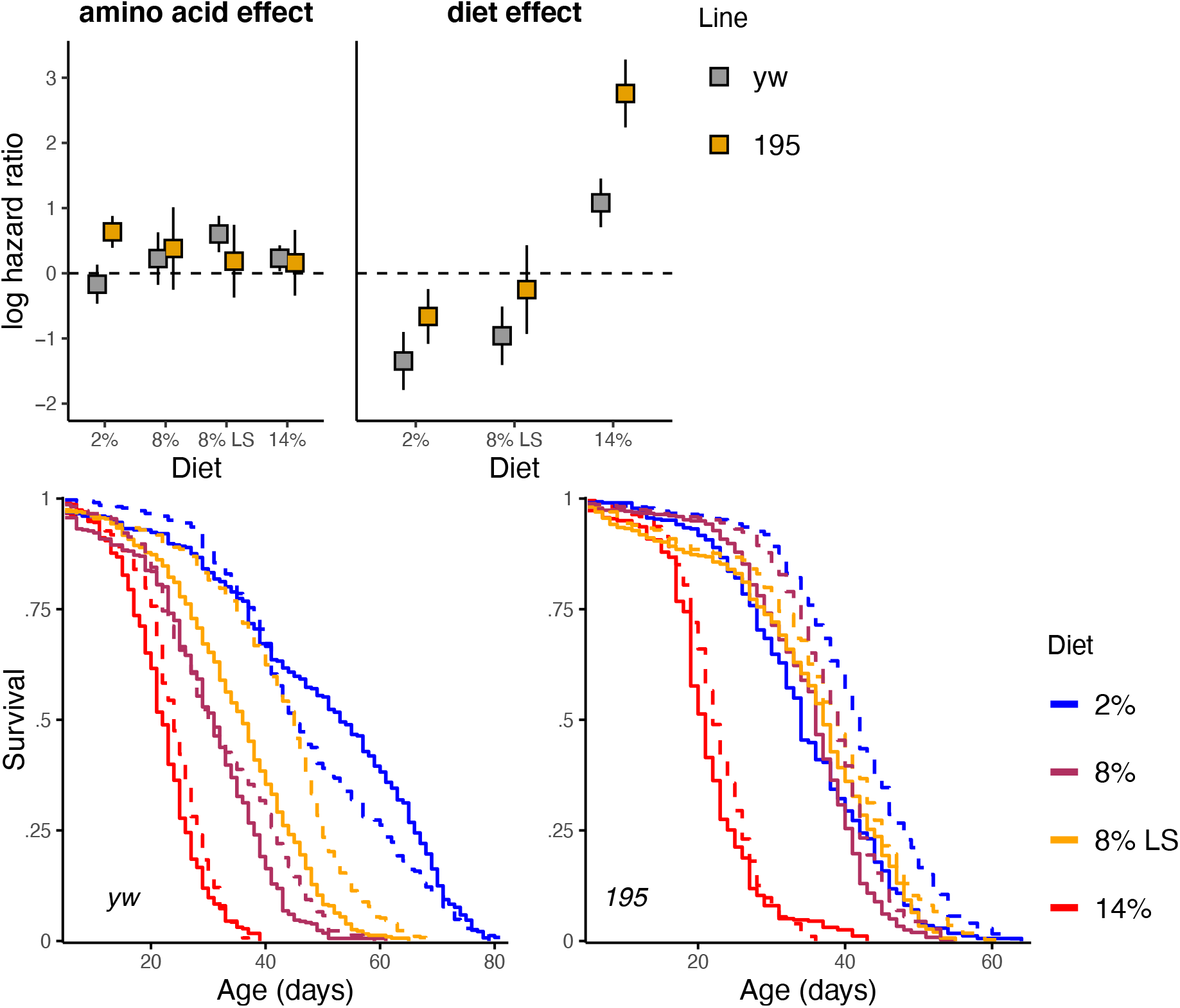
Top panel: Modulation of lifespan by EAA supplementation is small in comparison to diet, but varies by genotype. Log hazard ratios and their 95% CIs for both genotypes associated with amino-acid supplementation per diet (left, reference is control), and overall diet effects (right, reference 8%). Bottom panel: survival curves underlying the hazard estimates. Colour indicates the diets used with solid lines indicating amino acid supplementation, dash lines indicate control. Clearly EAA supplementation did not revert the lifespan gains achieved by modulating dietary yeast concentration.

Even though we used the exact concentration and composition of EAA as was previously shown to explain dietary restriction in the fly^5 replicated in 22^ a question could remain whether the supplementation had any substantial physiological impact in our experiments. The data collected on age-specific fecundity demonstrate, however, that egg laying is increased substantially with EAA supplementation and that this effect, in contrast to the effect on lifespan, is similar in magnitude to that of varying yeast concentration in the diet (Figure 2). Overall egg laying was increased in both genotypes to a similar degree by amino-acid supplementation (*yw*: 1.11 ± 0.19, P<0.0001;*195*: 0.89 ± 0.13, P=0.003, Figure 2), irrespective of the diet it was supplemented to (interaction diet by EAA addition, P > 0.16). For both genotypes analyses of the granularity of the patterns with age resulted in models including interactions of age with diet and with EAA supplementation, suggesting fecundity becomes differentially constrained with age depending on the nutritional environment (Table S2, Figure 2). Overall, these results suggest EAA supplementation fully rescues the loss of egg production when yeast concentration in the food is lowered, in contrast to the effects of yeast on lifespan.

**Figure 2.**
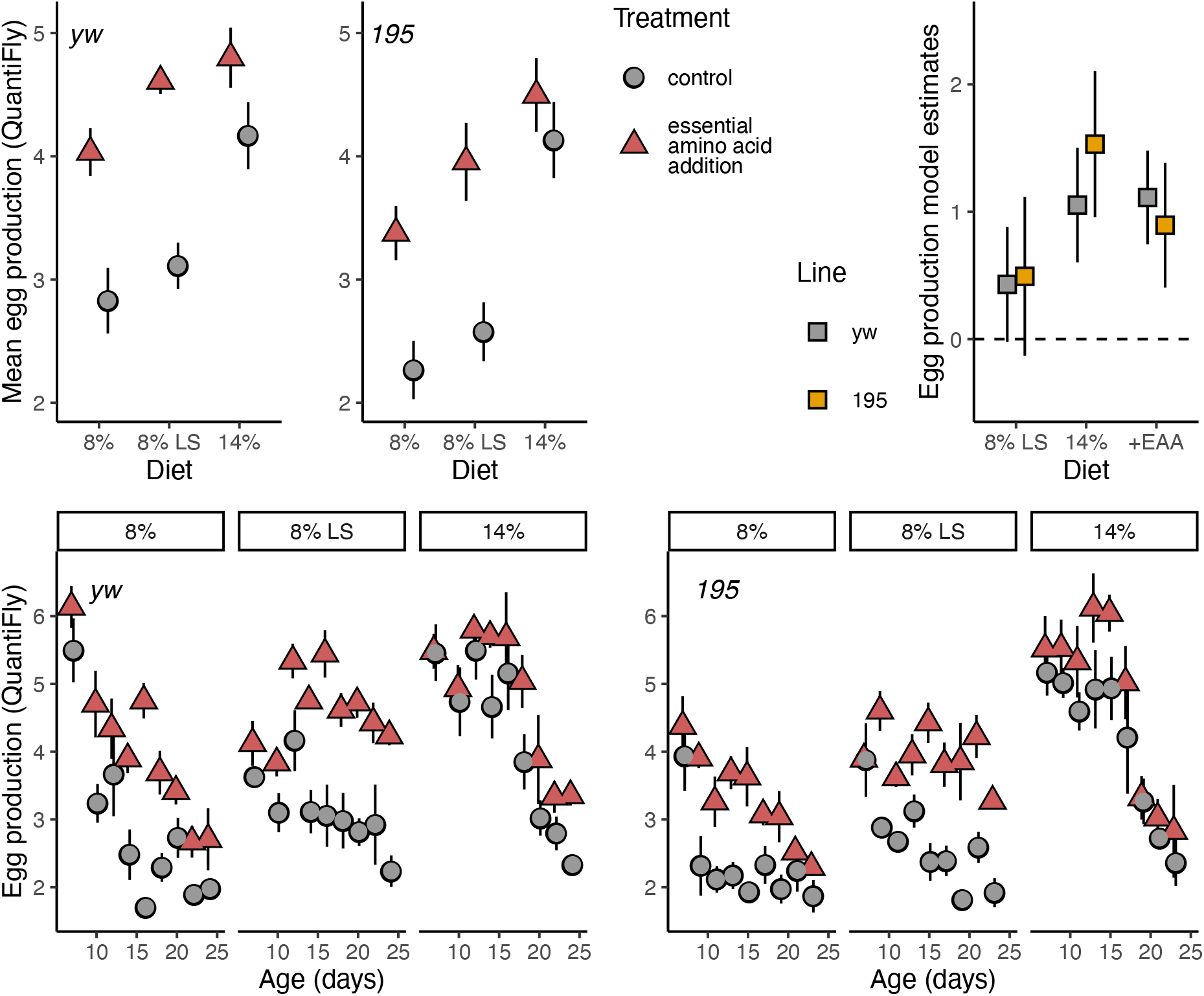
Left top panel shows mean egg production per fly (across 2 days) as measured using Quantifly. EAA supplementation increased egg laying on all diets and mimicked the fecundity gain from increased dietary yeast. Lower panels show age-specific fecundity plots per line per diet. Supplementation effects on fecundity vary by line, diet and age. Top right corner shows model estimates analogous to log hazard ratio plots, clearly showing a similar effect of EAA supplementation and increased yeast concentration (14%) on egg laying.

Interestingly, however, egg laying on the lowest yeast concentration (2%) was so low that counting via image analysis proved unreliable. Still, manual counting of eggs also revealed an increase in egg laying at this diet of lowest nutritional value when EAA were supplemented (Figure 3). This increase in egg laying was however modest and did not come near any of the egg production seen at the higher yeast diets (*yw*: 0.60 ± 0.10; *195*: 0.41 ± 0.05 at 2%, Figure 3, versus at 8%, Figure 2, *yw*: 2.88 ± 0.19; *195*: 2.37 ± 0.25). That egg production at low yeast conditions was not rescued by EAA supplementation suggests other nutrients limit egg production at this dietary condition. Arguably an even higher EAA dosage could increase egg laying at these diets. However, egg laying is reduced by 5-fold when yeast concentration is lowered, whereas the maximum estimated available EAA at a higher yeast concentration is only 3.1 times higher than the pure EAA supplemented (Table S3). Similarly, survival could reduce when a higher dose of EAA was supplemented, yet, the dietary modulation of log mortality hazard is 5.7 to 8.9 times that of EAA supplementation. Restriction of other nutrients than EAA therefore must explain a substantial part of the pro-longevity response of DR.

**Figure 3.**
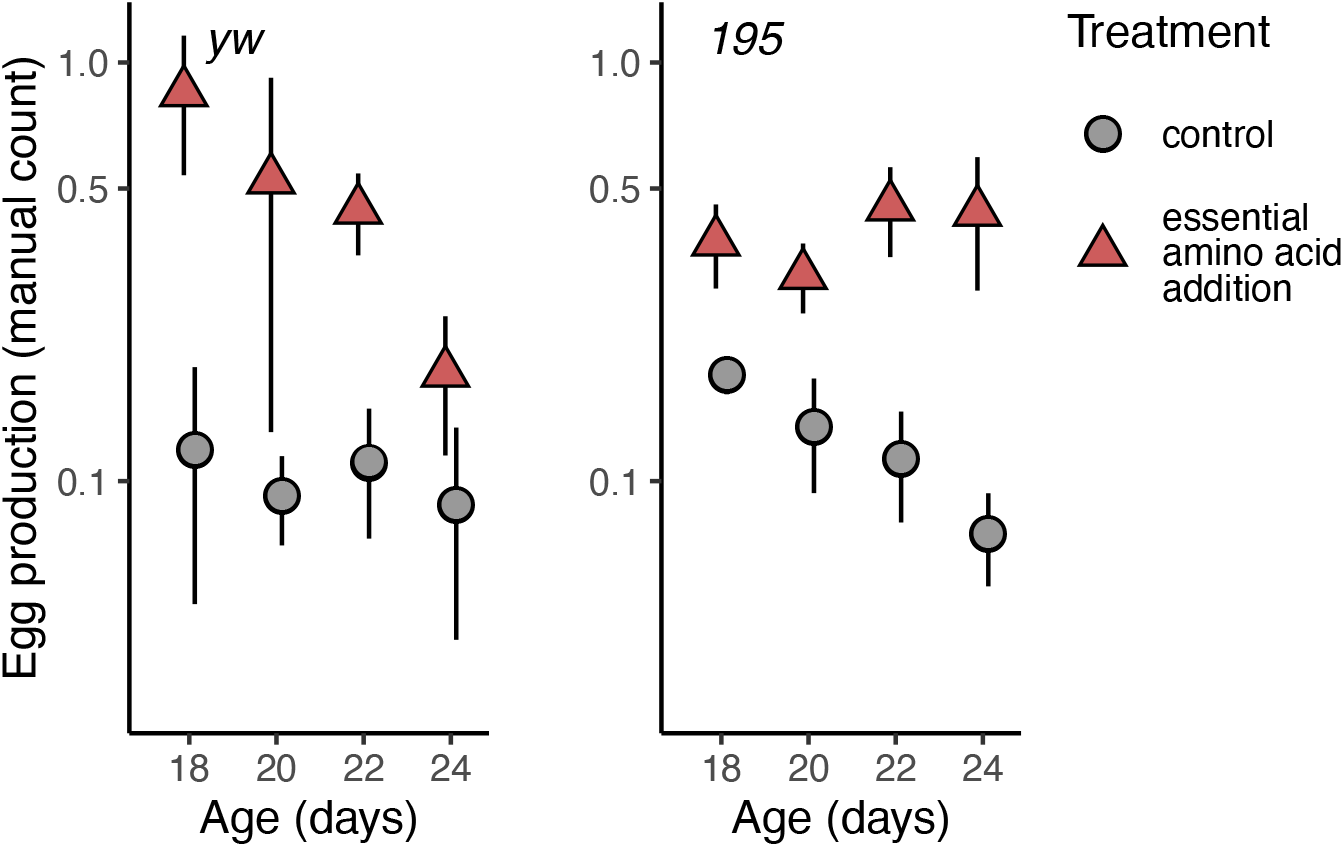
Mean egg production per fly on the lowest yeast diets (2%) from manual counts (across 2 days). EAA supplementation increased egg production but egg laying remains far lower than fecundity seen at the higher yeast diets (Figure 2), suggesting egg production is limited by other nutrients than EAA at this diet.

Carbohydrates (sugar) are currently considered to play a marginal role in determining longevity under DR, as dictated by the geometric framework^10,14^, but these studies are commonly only conducted on a single genotype^39,40^. We detect here a genotype specific effect of sugar on lifespan (Figure 1) that is interestingly accompanied by not a reduction but a non-significant increase in egg laying (Figure 2). Both genotypes tested also responded differently to amino acid supplementation in terms of lifespan, but similarly in terms of egg production (Figure 1–3). Genetic differences in the response to DR^41^ are of high interest, for translation^42^ and to understand the mechanisms of DR^43^. Until now however such differences have not suggested a differential response to specific nutrients. Our results suggest there is the potential for genetic variance in how nutrients shape lifespan. Potentially organisms should be viewed as a hierarchical set of physiological reaction norms^44^ to a range of nutrients. In this scenario DR becomes apparent when one of these reactions is limiting, and this need not be at the same level or the same nutrients for each genotype^39^.

## Conclusions

Previous work on the fly has been instrumental to the now consensus that dietary protein intake underlies the pro-health and pro-longevity benefits of DR across organisms, including our own species. Importantly, this idea has also shaped how amino acid availability might shape longevity through altered nutrient signalling. Our results now suggest that amino acid availability does not explain DR universally. These results are in line with recent experiments in mice that question the dominance of dietary protein in determining longevity, albeit with observable benefits to health^7^. Importantly, these insights now warrant a re-appreciation of how and in which circumstances specific nutrients, including amino acids, determine longevity and other key life history traits, such as reproduction. The metabolic networks that fuel bodily functions form a plastic network that shape phenotypic reactions to nutritional availability. The effects of nutrition on health are therefore probable to depend on genetic^39,43,45^, dietary^3,46^ and environmental^47,48^ interactions. Similar or differential pro-longevity physiology underlying DR could be triggered by differential nutritional interactions, and such could explain inconsistencies in our current understanding of DR and the mechanisms of ageing.

## Methods

Experiments used two genetic lines of Drosophila melanogaster - *195* from the DGRP^49^, and the yellow white (*yw*) lab strain. All flies were cultured on rich yeast media^40^ (8% autolysed yeast, 13% table sugar, 6% cornmeal, 1% agar, 0.225% nipagin and 0.4% propanoic acid). Cooked fly media was stored for up to two weeks at 4-6°C, and warmed to 25°C before use. Experimental diets consisted of 3% cornmeal, 1% agar and 0.225% nipagin, with dietary yeast and sugar varied to create the following experimental diets: low protein (2% yeast + 13% sugar), high protein (8% yeast + 13% sugar), high protein, low sugar (8% yeast + 5% sugar) and very high protein, low sugar (14% yeast + 5% sugar). The latter two diets are similar to the DR and fully-fed diets used in work that showed EAA are fundamental to DR^5^. Each of these diets was cooked with and without added essential amino acids, at the concentrations as in previous work^5,22^ (Table S3). Experiments were conducted in a climate-controlled environment with a 12:12 hour light-dark cycle, temperature at 25°C and 50-60% relative humidity.

### Longevity

Flies were grown in bottles on rich media and incubated at 25°C. In each of these bottles, 10 females (with 2 males) were allowed to lay eggs for 3 days. Bottles were given water daily if media appeared dry during larval development. When offspring began to eclose, individuals were transferred to mating bottles, where they were left to mate for ~48 hours. This was repeated every day until all flies had eclosed to generate age-matched cohorts. Flies were sorted under carbon dioxide anaesthesia (Flystuff Flowbuddy; <5L/min), and females were transferred to demography cages, specially designed to allow removal of deceased flies, and changing of food vial with minimal disturbance to living flies^40,50^. As in extreme conditions, and in some genotypes water availability can confound dietary effects^39^ all flies were supplemented with a vial of water agar ^as in 39^. Censusing was conducted every two days, with the food and water vial changed each time. Living flies which were stuck to the food vial or died as a result of sticking to the food or becoming trapped in any part of the cage, and flies which escaped, were right-censored.

### Fecundity

After 6 days of being on the experimental diets, vials from demography cages were photographed using a webcam under custom LED-lighting and images were analysed using QuantiFly^51^; machine learning software for automated image analysis for egg counting. Vials from cages on 2% yeast were counted manually using a dissection microscope, when it became apparent egg counts were too low to be reliably analysed with the QuantiFly setup.

### Statistical analysis

Lifespan data were analysed using time-to-event mixed-effects Cox proportional hazard models, with cage as a random term, implemented in ‘coxme’ in R^52^. Full models and comparisons with and without interaction terms between the dietary treatments (supplementation and diet coded as two separate independent variables) and line were used to test the main hypotheses. Estimates of individual models ran within each line and within each diet are presented as these will be less sensitive to deviations from proportionality of hazards and provide the best estimates of individual effects. Egg laying was analysed using linear mixed effects models in ‘lmer’^53^ in R, with cage as random term. Models on age-specific fecundity were simplified using backward selection, using the step function from ‘lmerTest’^54^. Egg counts were divided by the total flies in the cage at the time of fecundity measurement to correct for any differences in mortality, although uncorrected results yielded qualitatively similar results. In text, ± indicates standard error.

## Acknowledgements

We thank Laura Hartshorne and Gracie Adams for technical support. SLG is funded by the Faculty of Science, University of Sheffield. MJPS is supported by a Sir Henry Dale Fellowship (Wellcome and Royal Society; 216405/Z/19/Z) and an Academy of Medical Sciences Springboard Award (the Wellcome Trust, the Government Department of Business, Energy and Industrial Strategy (BEIS), the British Heart Foundation and Diabetes UK).

## Supplementary Tables

**Table S1.** Hazard ratio estimates of EAA supplementation (control as reference) and of diet (8% yeast as reference) from the cox proportional hazard mixed effect models. Estimates are from models ran within each genotype and or diet where applicable, as these estimates suffer least from any deviation from the non-proportionality assumption^40^. Higher log hazard ratios indicate a higher risk to die in the category stipulated.

|  |  | log HR | s.e. | p value | line |
| --- | --- | --- | --- | --- | --- |
| **Supplementation effect** |  |  |  |  |  |
| 2% | EAA | -0.17 | 0.15 | 0.27 | yw |
| 8% | EAA | 0.23 | 0.21 | 0.27 | yw |
| 8% LS | EAA | 0.60 | 0.14 | < 0.001 | yw |
| 14% | EAA | 0.23 | 0.10 | 0.02 | yw |
| 2% | EAA | 0.64 | 0.12 | < 0.001 | 195 |
| 8% | EAA | 0.38 | 0.32 | 0.24 | 195 |
| 8% LS | EAA | 0.19 | 0.28 | 0.51 | 195 |
| 14% | EAA | 0.16 | 0.26 | 0.53 | 195 |
| **Diet effect** |  |  |  |  |  |
| 2% |  | -1.35 | 0.23 | < 0.001 | yw |
| 8% LS |  | -0.96 | 0.23 | < 0.001 | yw |
| 14% |  | 1.08 | 0.19 | < 0.001 | yw |
| 2% |  | -0.66 | 0.21 | < 0.001 | 195 |
| 8% LS |  | -0.25 | 0.35 | 0.47 | 195 |
| 14% |  | 2.76 | 0.27 | < 0.001 | 195 |

**Table S2.** Model output following backward selection for age-specific egg laying (8%, 8% LS and 14% diets, Figure 2) within each genotype. For both lines the model including the diet by age and EAA by age interaction term proved to be the significantly better model, indicating EAA constrain reproduction differentially depending on the diet and age.

|  | Sum of Squares | Mean Square | Df numerator | Df denominator | F | p value |
| --- | --- | --- | --- | --- | --- | --- |
| **yw** |  |  |  |  |  |  |
| Diet | 7.54 | 3.77 | 2 | 20 | 10.60 | < 0.001 |
| Age | 110.54 | 13.82 | 8 | 160 | 38.84 | < 0.001 |
| EAA | 12.49 | 12.49 | 1 | 20 | 35.10 | < 0.001 |
| Diet * Age | 52.38 | 3.27 | 16 | 160 | 9.20 | < 0.001 |
| EAA * Age | 10.48 | 1.31 | 8 | 160 | 3.68 | < 0.001 |
| **195** |  |  |  |  |  |  |
| Diet | 14.54 | 7.27 | 2 | 20.15 | 24.76 | < 0.001 |
| Age | 79.05 | 9.88 | 8 | 154.28 | 33.65 | < 0.001 |
| EAA | 7.57 | 7.57 | 1 | 20.14 | 25.77 | < 0.001 |
| Diet * Age | 37.43 | 2.34 | 16 | 154.28 | 7.97 | < 0.001 |
| EAA * Age | 5.51 | 0.69 | 8 | 154.28 | 2.35 | 0.02 |

**Table S3.** Essential amino acid (EAA) concentration supplemented. Concentrations are as supplemented per litre of fly food and follow Grandison et al. 2009. The cooked fly food was split and a 20x stock solution dissolved in water was then added to cooled (~65 °C) media, the same volume in water was added to the control. The supplemented food and control food was therefore exactly the same apart from EAA addition. For comparison to EAA supplementation, the estimated EAA added through 6% yeast (2% to 8% to 14%) using two published estimates of EAA content (left^55^, right^56^).

| Essential amino acid | Concentration supplemented g/L fly food | Diet effect: estimated EAA added through 6% yeast g / L fly food |  |
| --- | --- | --- | --- |
| L-arginine | 0.43 | 1.01 | 3.02 |
| L-histidine | 0.21 | 0.4 | 0.52 |
| L-isoleucine | 0.34 | 0.88 | 1.15 |
| L-leucine | 0.48 | 1.37 | 1.66 |
| L-lysine | 0.52 | 1.73 | 1.43 |
| L-methionine | 0.1 | 0.16 | 0.47 |
| L-phenylalanine | 0.26 | 0.88 | 0.98 |
| L-threonine | 0.37 | 1.05 | 0.65 |
| L-tryptophan | 0.09 | 0.24 | NA |
| L-valine | 0.4 | 1.11 | 1.1 |
| *Total* | 3.2 | 8.84 | 10.9 |

